# Exploiting noise, non-linearity, and feedback for differential control of multiple synthetic cells with a single optogenetic input

**DOI:** 10.1101/2021.05.11.443599

**Authors:** Michael P May, Brian Munsky

## Abstract

Synthetic biology seeks to develop modular bio-circuits that combine to produce complex, controllable behaviors. These designs are often subject to noisy fluctuations and uncertainties, and most modern synthetic biology design processes have focused to create robust components to mitigate the noise of gene expression and reduce the heterogeneity of single-cell responses. However, deeper understanding of noise can achieve control goals that would otherwise be impossible. We explore how an “Optogenetic Maxwell Demon” could selectively amplify noise to control multiple cells using single-input-multiple-output (SIMO) feedback. Using data-constrained stochastic model simulations and theory, we show how an appropriately selected stochastic SIMO controller can drive multiple different cells to different user-specified configurations irrespective of initial condition. We explore how controllability depends on cells’ regulatory structures, the amount of information available to the controller, and the accuracy of the model used. Our results suggest that gene regulation noise, when combined with optogenetic feedback and non-linear biochemical auto-regulation, can achieve synergy to enable precise control of complex stochastic processes.

## Introduction

Synthetic biology seeks to develop and characterize biological circuits and modular components that can be reliably re-engineered, re-assembled, and controlled to produce complex biological behaviors. ^1^ The design and implementation of modular components as simple functional units^2^ capable of being assembled to perform desired regulatory needs has yielded powerful capabilities of synthetic cells to perform specific actions in response to specific stimuli. ^3^ Early advances led to the development of programmed cells that are capable of complex logic like switching, self-regulation, and fast acting control. ^4^ Genetically engineered switches were originally built in bacteria, ^5^ but have been extended to yeast, ^6^ mammalian cells, ^7^ and even multicellular plants. ^8^ In turn, these simple switches have led to more complex engineered biological systems and biotechnologies capable of performing tasks like controlling cells to behave as digital displays. ^9^

Much of the design process for synthetic biology has focused directly on building better biological components, such as creating more sophisticated gene regulatory structures, ^10–12^ introducing new response elements or reporters, ^13^ or introducing more orthogonal cellular signals. ^14^ Development work on gene regulatory structure has led to substantial advances in phenotype control in plant biology and targeted or modified protein turnover in therapeutics. ^15,16^ Improvements to response reporters and the experimental techniques used to analyze cells, especially in the form of fluorescent protein reporters, ^17,18^ real time single-gene MCP-MS2-based transcription elongation assays, ^19^ and nascent chain translation elongation assays^13,20–22^ has made it possible for cells to transmit their internal states to human observers. These technologies allow for more observation and control, not just at the protein level, but at the gene and RNA levels as well. Advances in cellular signals have also introduced the potential for synthetic regulatory modules in separate cells to communicate with one another and control multicellular population dynamics. ^23–25^ For example, by considering a simple model for the effect of cellular quorum sensing on cell densities, simple circuits can be tuned to control cell densities. ^26^

Although the above advances have been developed primarily to control autonomous biological behaviors, these improvements to regulatory structures, response reporters, and signaling capabilities can also provide a framework to allow observers or external electronic devices to monitor cellular environments and dynamically reprogram the cellular logic. When coupled to advances in microfluidics, these capabilities introduce a new part-biology-part-machine (or cyber-organic^27^) paradigm that adds new possibilities for distributed external and internal biological control of synthetic biological circuits. ^28^ In particular, recent developments in optogenetics^29,30^ have greatly increased the speed and sensitivity by which external signals can be communicated from humans or machines to cells. Using these advances in microfluidics and light-activated gene regulatory elements is rapidly improving the potential to integrate carbon- and silicon-based circuits, which in turn is making hybrid bio-electronic circuits far more powerful than before.

A key challenge to integrating cell-based genetic circuity with electronic control is that an uncountable number chemical species and regulating bodies must diffuse and interact with one another in space and time within each cell. Tractable analysis requires immense simplifications of these infinitely complex and chaotic dynamics, and such simplifications naturally result in large uncertainties that can only be accounted for through the introduction of approximate models with stochastic analyses. One of the greatest challenges to improving externally controlled cellular behaviors is that this ‘noise’ in gene regulation introduces large amounts of single-cell heterogeneity which must accounted for. ^31–33^ Under the context of this noise, cellular responses are probabilistic–their distributions may shift gradually and even be statistically predictable under environmental or genetic manipulations, ^34–36^ but individual cells appear to behave at random with very little information about their instantaneous external environment.^37^ Unfortunately, this noise makes it difficult to precisely predict how or when an individual cell will respond to a new environmental stimulus. Although recent work has shown that external feedback can control and reduce cellular heterogeneity within a large population, ^38^ it may seem unlikely that any feedback control strategy could reliably guarantee that specific cells within a population will respond as desired, and independently of their initial conditions.

Most efforts on external feedback control in synthetic biology have focused on the use of chemical or optical inputs to manipulate cell population averages ^39^ or to control individual cells within a larger population. ^39^ Such efforts can be classified as single-input-single-output (SISO) or multiple-input-single-output (MISO) control in that they seek only to push cells to a single phenotype. For example, recent experimental and computational studies ^40–42^ have used periodic chemical input fluctuations to control one cell or a population of multiple cells to be as close as possible to the same unstable fixed point, a task that is similar to the inverted-pendulum problem in classical control theory. Other recent work has sought to control multiple individual cells, each with their own tailored optogenetic inputs, ^29^ which corresponds to multi-input-multi-output control (MIMO). SISO and MISO control are limited to control only a single cellular response at a time, while MIMO requires advanced hardware such as digital micro-mirror devices that devote a separate input to each individual cell. ^29,43^ However, the combination of synthetic biological designs, precise external controls, and quantitative measurements and models of single-cell noise could create new opportunities for single-input-multiple-output (SIMO) control, where multiple individual cells could be controlled to achieve *different phenotypes*, but requiring only a single input signal. In this article, we explore the potential of a noise-enabled SIMO control strategy that is similar to a hypothetical Maxwell’s Demon (MD), who watches the random process and identifies short instances in time when specific cells have randomly increased sensitivities to small perturbations. ^44^ In the context of synthetic biology, we explore how advances in fast fluorescent reporters allow this MD to watch biological responses; how noise breaks the symmetry between identical cells in identical environments; and how advances in optogenetics may enable the MD to drive specific cells toward specific phenotypes, even when each cell always experiences the exact same input signal as every other cell.

Through simulation of multiple models that have been parametrized from existing bulk level optogenetic control experiments, we demonstrate that realizing an optogenetic MD for use in genetic regulation applications requires not only careful consideration of the cellular regulatory systems to be controlled, the fluorescent sensors to be observed, and the optogenetic inputs to be delivered, but it is also necessary to build predictive stochastic models and deterministic SIMO control algorithms to serve as its brain. In this article, we use a combination of simulations and theoretical analyses to explore how existing biological parts could be combined with models in the context of optogenetics to control the gene expression of multiple cells at once, even when both cells receive the same light signal at the same time and have the exact same genetics. We then introduce a new probabilistic model predictive control (pMPC) strategy that can in principle control multiple cells to different phenotypes, even when only observing a single cell. Finally, we show that controls that are designed and optimized using one approximate model of the biological system can be effective to control a much more complex biological process whose mechanisms are not considered during the controller design.

## Results and Discussion

We start by defining a set of two models, each with a different level of complexity, to describe the dynamics of optogenetically activated T7 polymerase in temporally-varying light conditions. We then use experimental data from Baumschlager et al^45^ to independently constrain each of these models to the same data. We then take the simpler of the two models and extend it to include a typical auto-regulation module, and we examine the performance of this extended model under different fluctuating input signals at different frequencies using deterministic and stochastic analyses. We then show how intrinsic noise in the system dynamics can be utilized by a feedback controller to break symmetry in the process and force a system of two cells each to obtain specified phenotypes and independent of initial conditions. We then propose a new probabilistic model predictive controller scheme that is capable to differentially control multiple cells even when only one is directly observed. Finally, we demonstrate in principle that an optogenetic controller that is identified using a coarse-grained simplified model is capable to control behavior of a more complicated system with different and hidden dynamics.

### A data-constrained model for light-induced T7 polymerase activation and gene expression

We begin by developing an *unregulated* model (denoted as ℳ_*U*_) to describe the light-induced activation of an optogenetically controlled T7 polymerase as studied in Baum-schlager et al. ^45^ As depicted in Fig. 1A, this system contains two light activated T7 domains (*T*7_*n*_ and *T*7_*c*_) that are produced at constant rates *k*_*n*_ and *k*_*c*_, and which degrade in a first order decay process with rate *γ*_*M*_. These domains dimerize when subject to light activation leading them to form an active T7 polymerase at a light dependent rate of *u*(*ϕ*) and the complex dissociates at a rate of *k*_*i*1_ and decays at a rate *γ*_*T*_. The T7 polymerase dimer can bind to, or unbind from, the gene at rates of *k*_*f*2_ and *k*_*i*2_, respectively. When bound, active protein production occurs at a rate of *k*_*f*1_, where transcription and translation are lumped into a single event. Proteins are assumed to degrade according a first order rate process with rate *γ*_*P*_. These interactions are described by the following reactions

**Figure 1:**
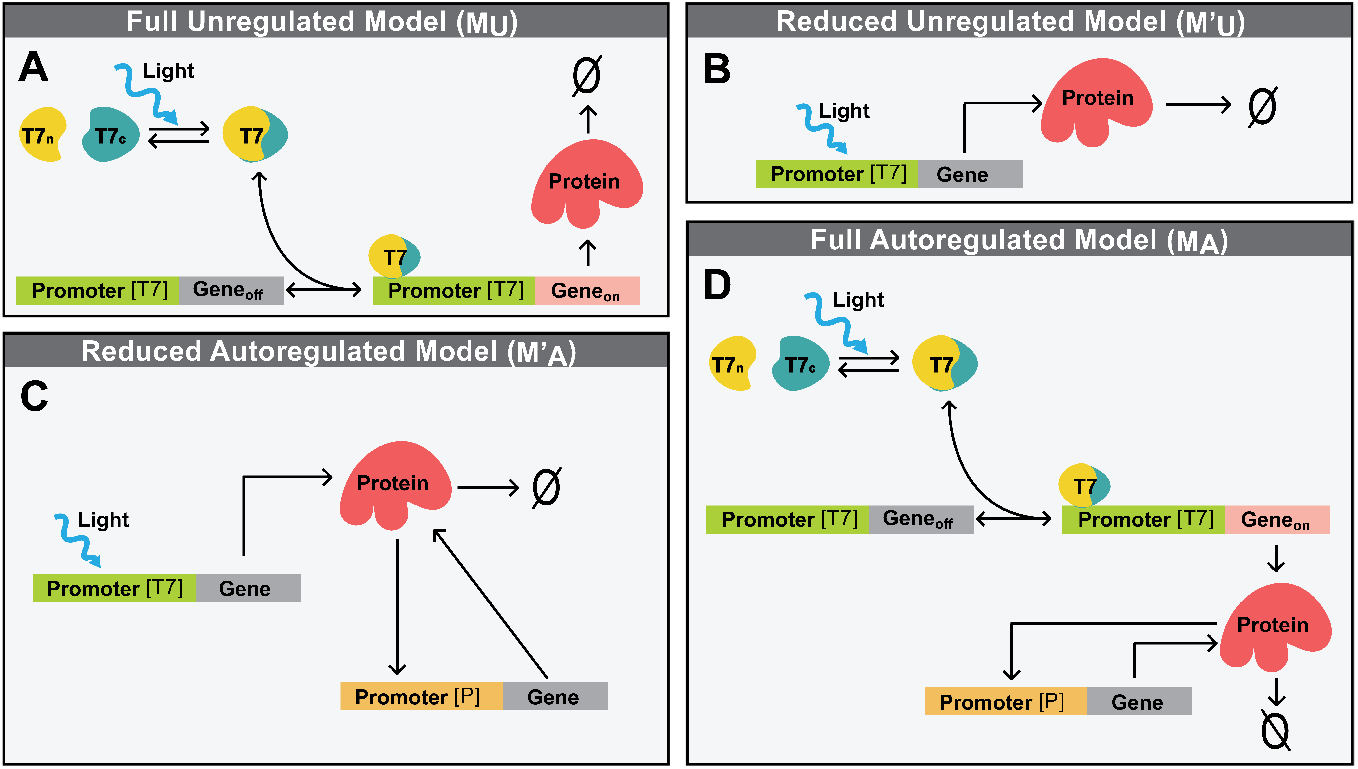
Diagrams of models for light-induced gene regulation with and without auto-regulation. (A) The full physical model (ℳ_*U*_) describes the full mechanistic processes of T7 polymerase dimerization and gene activation in addition to protein production and degradation. (B) A simplified model 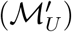 lumps together all T7 dimerization and gene activation dynamics so that protein is produced at a light-controlled rate. (C) The simplified model is extended 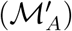 to include auto-regulation through the addition of a secondary promoter. (D) The full mechanistic model is also extended (ℳ_*A*_) through inclusion of the same auto-regulation promoter.

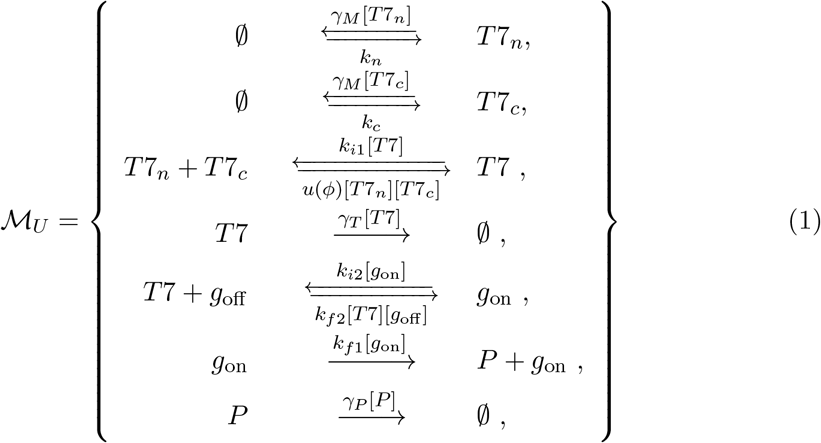

where the first two bidirectional reactions describe production and decay of *T*7_*n*_ and *T*7_*c*_; the third bidirectional reaction describes light-induced reversible dimerization, where the light induction level is denoted as *ϕ* and its effect is modeled by the function *u*(*ϕ*); the fourth unidirectional reaction describes the decay of the T7 dimer; the fifth bidirectional reaction describes the T7 association and dissociation to the gene; and final two unidirectional reactions describes the production and degradation for the resulting protein product, where transcription and translation have been lumped into a single reaction. The rate for each reaction is given directly above or below its respective arrow.

The ODE for ℳ_*U*_ can be written in vector form as

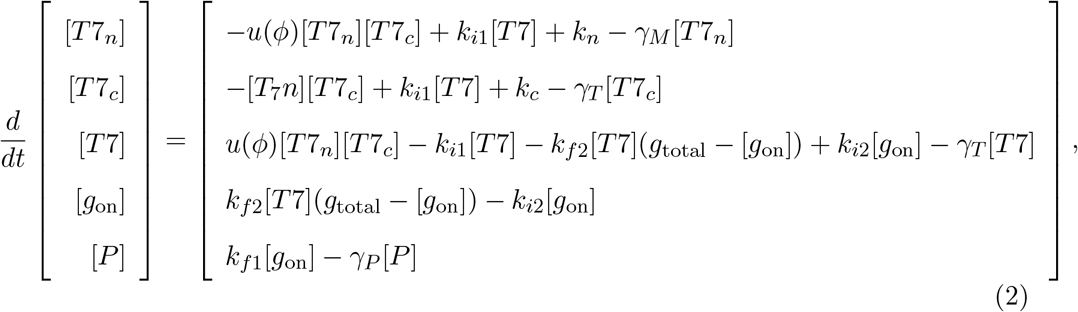

where we have assumed a fixed number of gene copies [*g*_total_] = [*g*_on_] + [*g*_off_] to remove the variable [*g*_off_] from the equations. Once written in this form, Model ℳ_*U*_ can be integrated numerically for any given set of initial conditions and parameters. The model parameters were then fit to the experimental data from Baumschlager et al, ^45^ in which the system was subjected to three different levels of UV radiation:

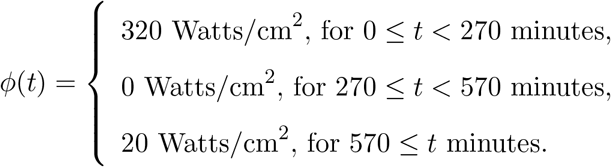

The parameters of ℳ_*U*_ and {*u*(*ϕ*_1_), *u*(*ϕ*_2_), *u*(*ϕ*_3_)} are simultaneously fit to the measured time series fluorescent protein trajectory. This fit suggests *u*=[0.4060, 0.00, 0.0400] molecules^−1^min^−1^ when *ϕ*=[320, 0, 20] Watts/cm^2^, respectively. To interpolate for intermediate values of light intensity, the function *u*(*ϕ*) is then defined as the cubic spline of the three data points and is shown with a red line in Fig. 2B. The resulting fit of the model to the data is shown by the red dashed lines in Fig. 2A, and the remaining parameters of the model fit are shown in Table 1.

**Table 1:** Parameters of the full model system.

| Parameter Name | Parameter Value | Units |
| --- | --- | --- |
| $k_n$ | $2.00 \times 10^{-1}$ | molecules/min |
| $k_c$ | $6.00 \times 10^{-1}$ | molecules/min |
| $k_{i1}$ | $2.00 \times 10^2$ | $\text{min}^{-1}$ |
| $\gamma_M$ | $5.00 \times 10^{-2}$ | $\text{min}^{-1}$ |
| $\gamma_T$ | $5.00 \times 10^{-2}$ | $\text{min}^{-1}$ |
| $k_{i2}$ | $5.00 \times 10^{-1}$ | $\text{min}^{-1}$ |
| $k_{f2}$ | $5.00 \times 10^{-1}$ | molecules/min |
| $k_{f1}$ | $1.42 \times 10^{-2}$ | $\text{min}^{-1}$ |
| $\gamma_P$ | $2.03 \times 10^{-2}$ | $\text{min}^{-1}$ |

**Figure 2:**
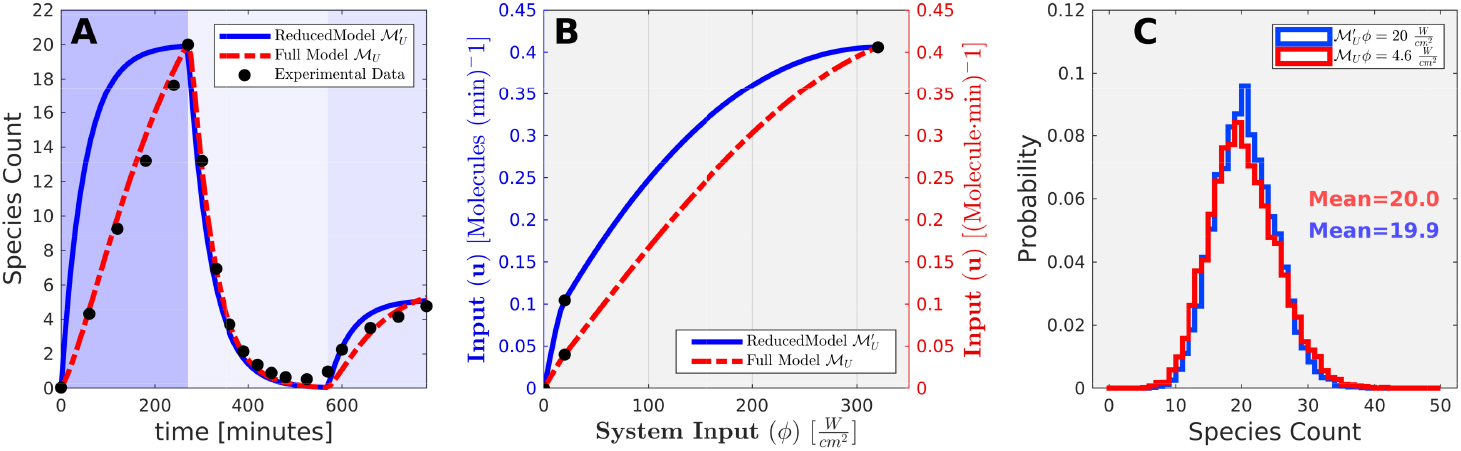
Parameter estimation of full and reduced unregulated models, ℳ_*U*_ and 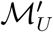, respectively. (A) Fits of ℳ_*U*_ (red line) and 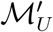 (blue line) to experimental data from Baumschlager et al.^45^ (black dots).(B) Calibration curves representing the conversion of light [Watts per cm^2^] to the associated reaction rates for ℳ_*U*_ (red) and 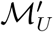 (blue). (C) Steady state histograms predicted by models ℳ_*U*_ (red) and 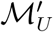 at inputs that are calibrated to results in an average expression of 20 molecules per cell.

Having determined a baseline ODE-based model that yields a good fit to existing experimental data for the system’s temporal response, in the next sections we will specify a much simpler, but less accurate, version of this model and use that approximate model to perform stochastic analyses, suggest design modifications, and specify a controller that can drive differential gene expression among two or more cells using a single external input signal.

### A substantially reduced and approximate model for T7 activation and gene expression

Although we were able to find many good parameter sets so that model ℳ_*U*_ could match the data, ^45^ identification of a single unique ‘best’ parameter set is infeasible due to the high number of parameters, severe limitations on available experimental data, and sloppiness ^46^ in the model parameters. Finding a simpler, but better constrained model would not only help to reduce sensitivity to unknown parameters, but could also dramatically reduce computational costs when using that model for design decisions or for the specification of feedback control strategies. To simplify these model reactions and to obtain a more identifiable parameter set, we next propose a simple generalized birth-death model in which the protein production rate is given by *k*_0_+*u*′(*ϕ*), where *k*_0_ is the baseline production rate with no light input, and *u*′(*ϕ*) is a light-dependent control input function. We use the apostrophe notation (·)′ to denote use of the reduced model in which the units of the control signal at a given light intensity have been adjusted to molecules per minute. Under this simple rule and assuming linear decay at rate *γ*_*P*_, the approximate dynamics of *P*(*t*) are written simply as:

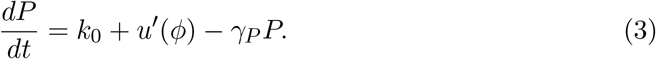

In this model, which we denote as 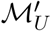, the parameters *k*_0_ and *γ*_*P*_ and the specific values of *u*′(*ϕ*) at *ϕ* ∈ {320, 0, 20} Watts/cm^2^ are again simultaneously fit to capture the time dynamics of the experimental data. ^45^ This fit suggests *u*′(*ϕ*)=[0.4060, 0.00, 0.1044] molecules/min when *ϕ*=[320, 0, 20] Watts/cm^2^ respectively, and the cubic spline of these three data points yields the calibration curve *u*′(*ϕ*) as shown in Fig. 2B (blue line). The resulting fit of model 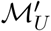 is shown in the solid blue line Fig. 2A, and the remaining parameters for the reduced model are shown in Table 2.

**Table 2:** Parameters of the reduced model system.

| Parameter Name | Parameter Value | units |
| --- | --- | --- |
| $\gamma_P$ | $2.03 \times 10^{-2}$ | 1/min |
| $k_0$ | $1.00 \times 10^{-4}$ | molecules/min |

We next extended both the original model and the simplified model to include discrete stochastic events for protein production and degradation, as well as for T7 dynamics for the full model. Using the exact same parameter values as for the ODE analysis, we then simulated the model using the Stochastic Simulation Algorithm (SSA, ^47^). Fig. 2C shows the probability distribution of *P* sampled using 10^5^ independent SSA runs, each simulated to 3000 minutes and only using the last data point of each simulation for either the full model (red) or the simplified model (blue). Both models provide reasonably good, but not identical, matches to the mean protein levels and the overall time scales as shown in Fig. 2A, and we observed strong agreement in their stationary distributions. However, it remains to be seen if the differences in architecture and time scales need to be accounted for when we use the simplified model 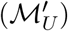 to guide our modification of the gene regulatory system and to design a feedback controller that remains effective even when applied to the more complex original model (ℳ_*U*_).

### Addition of an autoregulation motif creates light-dependent bistability

Starting from the simplified model,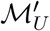, we next asked if a common auto-regulation motif could be added to introduce light-dependent bistability for the system and so that we could explore how that added motif would impact the controllability of the overall system. Fig. 1C shows a schematic of the new model denoted as 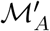, which has auto-regulation due to the addition of a secondary promoter that is self-activated by protein *P* to produce more of itself. To incorporate this auto-regulation behavior, a Hill function production rate is assumed with high cooperativity, and the new rate equation becomes:

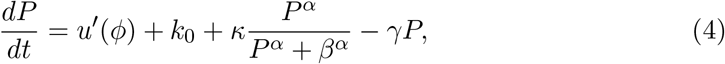

where the first term *u*′(*ϕ*) corresponds to the control input as a function of light input; the second and third terms correspond to the Hill function activity of the autoregulation promoter with leakiness *k*_0_, and the final term corresponds to the first order decay of the protein. The Hill parameters (*β, α*) of 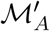 are chosen in order to exhibit bi-modal behavior in the dynamics of *P*. All parameters of this auto-regulatory model are shown in Table 3.

**Table 3:** Hill function parameters for auto-regulation motif.

| Parameter Name | Parameter Value | Units |
| --- | --- | --- |
| $\kappa$ | $4.06 \times 10^{-1}$ | molecules/min |
| $\gamma$ | $2.03 \times 10^{-2}$ | min<sup>-1</sup> |
| $\beta$ | 20 | molecules |
| $\alpha$ | 8 | unitless |
| $k_0$ | $1 \times 10^{-4}$ | molecules/min |

Fig. 3A shows bifurcation diagrams for the simplified unregulated and auto-regulated models, 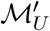 and 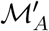, respectively. These diagrams show that as the light input sweeps *slowly* from zero to 0.5 Watts per cm^2^, bifurcation and hysteresis become apparent in the auto-regulated model (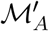, light/dark blue), but these effects are not observed in the unregulated model (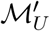, orange/red). For either model, low and high light inputs each result in a single stable point at low or high expression, respectively. For intermediate light inputs, however, two history-dependent stable points coexist for 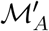, and it is possible for two cells to maintain different stable points (or phenotypes) provided that the light intensity remains in the bi-stable region and that the cells begin in the separate basins of attraction for the different stable points.

**Figure 3:**
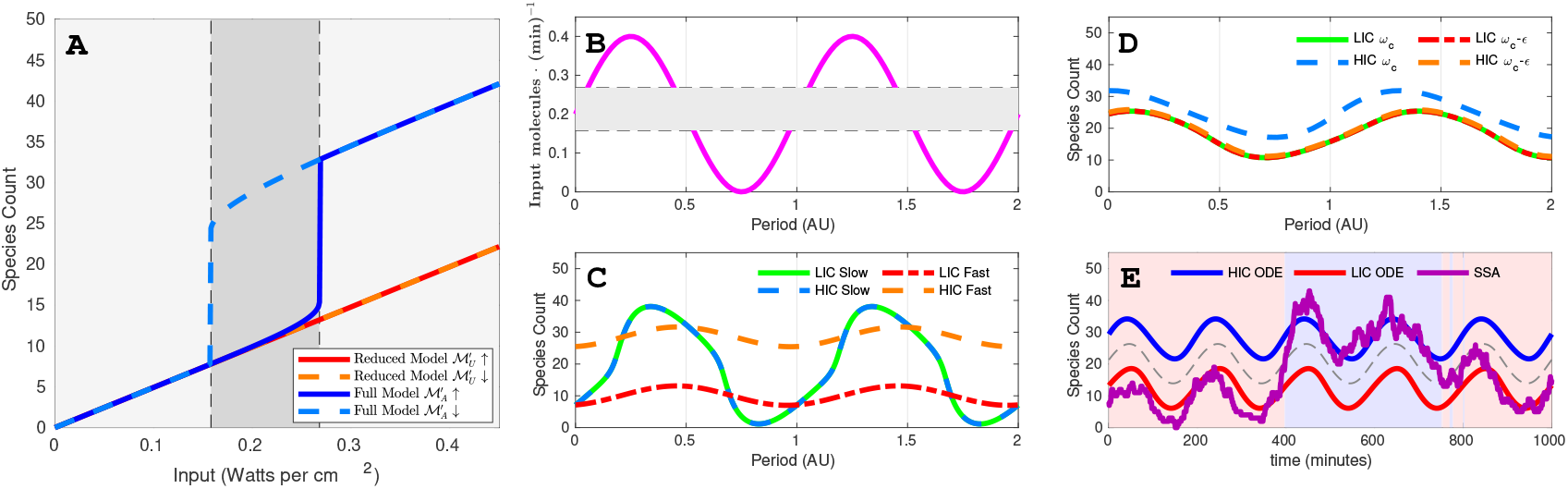
Input-output analysis for the simplified model with and without auto-regulation (Models 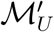 and 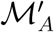). (A) Bifurcation (hysteresis) analyses of 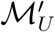 (red shades) and 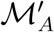 (blue shades). Solid (dashed) lines depict the change in the steady state solution as UV intensity is increased (decreased). Bi-modality (i.e., two possible steady states at the same control input) and hysteresis (i.e., different paths of solid and dashed lines) only occur for 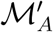. (B) Sinusoidal input used for temporal excitation shown for two periods. (C) Steady state temporal behavior of 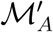 under sinusoidal input shown for two periods. Dashed lines correspond to high initial conditions, and solid lines correspond to low initial conditions. Different colors correspond to different input signal frequencies: fast (0.01 RPM, red/orange), slow (0.001 RPM, blue shades). All plots show steady state temporal response after a transient time of at least two oscillation periods or 3500 minutes, whichever is longer. (D) Same as (C) but with frequencies 0.0043522 RPM (red/orange) and 0.0043525 RPM (blue shades). The 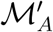 maintains memory of initial condition provided that input frequency is greater than a critical value (i.e., solid and dashed lines remain distinct). (E) Capability of Model 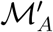 to track inputs assuming stochastic fluctuations as analyzed using an extended SSA^47^ with extra reactions. ^48^

In the hysteresis plots of Fig. 3A, it is assumed that the light input sweeps very slowly so that the response reaches equilibrium at each light level before subsequent changes. However, in the context of feedback control, we are more interested in how cells respond to light fluctuations at faster, transient time scales. Therefore, we next test the stability of input-to-output behaviors under time-varying inputs. We start simulations for two cells with identical parameters, but at different initial conditions (i.e., one at a high initial concentration of 40 molecules per cell and another at a low concentration of 0 molecules per cell), and we subject these both to the same sinusoidally-varying input signal whose range encompasses both bifurcation points as shown in Fig. 3B. Fig. 3C shows the steady state trajectories for two pairs of such systems, where dashed lines correspond to trajectories that start at low initial conditions and solid lines correspond to trajectories starting at high initial conditions. In all cases, the system is subject to at least four cycles or 3500 minutes so that transient dynamics have had time to decay, and the time axis is scaled to show the response over two oscillation periods. Fig. 3C (blue shades) shows that when the input frequency is slow (*ω*_*slow*_ = 0.001 rotations per minute (RPM)), the system loses memory of its initial condition and the trajectories from both initial conditions decay to a single trajectory. However, when the frequency is fast (red and orange trajectories, *ω*_*fast*_ = 0.01*RPM*), the system can maintain memory indefinitely. Fig. 3D shows that the cut off frequency for this maintenance of memory is sharp in that memory is possible at a frequency of *ω*_*c*_ (blue shades) but is lost at a slightly slower frequency of *ω*_*c*_ − *ε*, where *ω*_*c*_ = 0.0043525 (red/orange) RPM is the critical frequency and *ε* = 3 × 10^−7^ RPM is a small perturbation to that frequency.

### Intrinsic noise can drive cells to switch phenotypes

Using deterministic analyses of the bistable model, we have seen that two cells with different initial conditions maintain separate phenotypes as they respond to the same fluctuating input signal. The flip side of this deterministic result is that two fully converged solutions of the same ODE never diverge, such that two cells starting at the exact same initial condition will never express unique phenotypes even when bi-stability is possible. However, low copy numbers of important regulatory molecules (DNA, RNA and proteins) often result in stochastic fluctuations in cellular concentrations (also known as ‘intrinsic noise’) that dramatically affects both of these results. When added to a bistable deterministic process, noise can drive two cells starting at the same initial phenotype to diverge or even drive two cells to exchange phenotypes by chance over time. ^49,50^ With this possibility in mind, we next ask how noise would affect the ability of cells to track a temporally varying input signal. For this, we examined the production and degradation reactions and converted model ℳ_*A*_ to an equivalent discrete stochastic model with the exact same rate parameters, and we explored how discrete intrinsic noise of stochastic models could be used drive cells to separate phenotypes. To extend the SSA to approximate time-varying inputs, we adopted an approach similar to that in Voliotis et al, ^48^ and added a fast ‘null event’ reaction that updates the clock and input signal value on a time scale that is much faster (i.e., average of 100 events per period of the input signal) than that of the input signal fluctuations. We then compared the ODE and the SSA analysis of 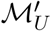 under a sinusoidal input with moderate input frequency of *ω* = 5.00 × 10^−3^ RPM *> ω*_*c*_ for which the ODE trajectories maintain memory of their initial conditions. Although the ODE solutions (smooth lines in Fig. 3E) will never converge, the stochastic trajectories (purple fluctuating trajectories) switch occasionally between the two fluctuating phenotypes. In other words, with the addition of noise to the system, each cell can slowly ‘forget’ its original configuration. Moreover, the probability of switching depends on the transient stochastic state of the process and the frequency and amplitude of external input fluctuations. Previous studies have observed similar effects for how noise creates variation in a population of cells, and past feedback control efforts have sought to counteract this variation to keep all cells at a chosen (and in some cases unstable) phenotype. ^40–42^ In the next section, we do not try to reduce variability among cells, but rather we seek to exploit the condition- and time-dependent disruption of symmetry to push one cell to a chosen phenotype while forcing another cell or group of cells toward a different chosen cell fate.

### Finite State Projection analyses uncover effective strategies to differentially control two cells using a single input

We consider a system of *N*_*c*_ cells, each with identical regulatory mechanisms and parameters, but whose fluctuating protein concentrations at time *t* are denoted by 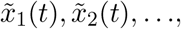, which we can arrange into the vector 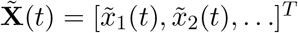. Here, the notation 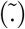 denotes that the corresponding quantity (e.g., protein copy number) is the result of a stochastically fluctuating process. Our overall goal is to design a feedback control law to force 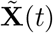 as close as possible toward an arbitrary target state 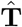. For an example with two cells, 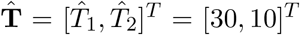 would correspond to having the first cell in the high expression phenotype and the second cell in the low expression phenotype. In general, this fluctuating control signal could depend on measurements of 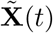 and/or the current time according to some as yet to be determined control function 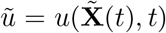. We define a cost function as the expected squared Euclidean distance between 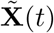 and 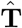 at steady-state, which can be written as

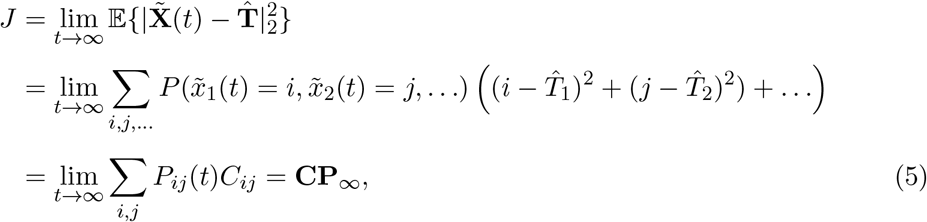

where **C** is a constant vector of squared Euclidean distances of each state from the target, and **P**_∞_ is the steady-state probability mass vector (i.e., the stationary probability for each unique value of 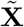).

As described in Methods, the master equation for a finite number of cells subject to a fluctuating state- or time-dependent input signal can be written as

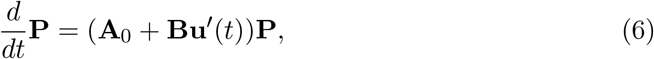

where 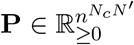 is the probability mass vector for all possible states; 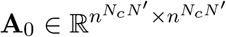 is an uncoupled infinitesimal generator; 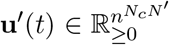 is a (potentially time varying) vector of control inputs with one entry for every distinct state in the system’s state space; and **B** is a fixed tensor that operates on **u**′ to adjust the master equation to account for the optogenetic input. Explicit examples for the construction of quantities **A** and **Bu**′ for different control laws are provided in Methods.

At first, we consider the special case where the control signal depends only on the current state at each instant in time. In this case, the vector **u**′ is constant with respect to time and depends only on the enumeration of the possible states. As such, the infinitesimal generator in Eq. (6) reduces to a time-homogeneous master equation for a standard discrete state Markov process. We note that the control signal *ũ*′(*t*) may still fluctuate due to changes in 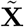 and can be written using the notation 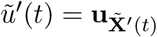 to denote that the index of the vector **u**′ is specified by the instantaneous state 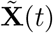.

By changing the specification of **u**′, we can further simplify this class to consider several different types of state-based control rules. Here we consider three different possibilities: an UnAware Controller (UAC) that has no information about any cells to make its control decision:

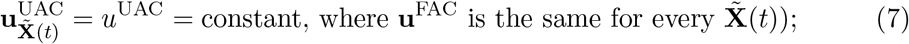

a Fully Aware Controller (FAC) which has complete knowledge of all cells 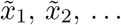 and therefore uses the full state vector to make its control decisions:

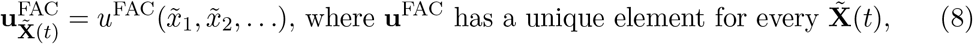

and a Partially Aware Controller (PAC) which uses only information of a single cell, e.g., *x*_1_, to make its control decisions:

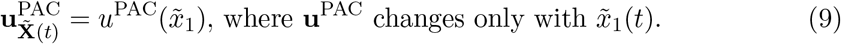

For our specific case of model 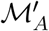, the FSP truncation size is *n* = 50, and the number of species is *N*′ = 1; thus for two cells (*N*_*c*_ = 2), the matrix **A**_0_ ∈ ℝ^2500*×*2500^.

With the formal definition of the cost function *J* from Eq. (5) and the CME from Eq. (6), we can then compute the gradient of the cost function with respect to the control law as (see derivation in SI):

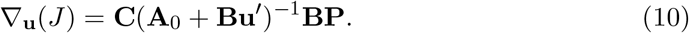

Having specified the cost function gradient with respect to the controller, we run an optimization algorithm (see Methods) to search along the gradient to find local minima for the cost function. We note again that for each of these optogenetic Maxwell’s Demon control strategies, every cell experiences equal inputs at every instant in time, although the magnitude of that single input changes in time as the cells fluctuate to different states.

Starting with the unaware (UAC) and fully aware (FAC) controllers, we used the FSP approach and local optimization (see SI for details on the optimization procedure) to find a locally optimal control strategy in the form of the constant *u*^*UAC*^ or the two-dimensional scalar field *u*^*FAC*^(*x*_1_, *x*_2_), and with and without the addition of auto-regulation to the model. The resulting optimal controllers are presented in the top row of Fig. 4, where the color at each point represents the control input magnitude *u*^*xAC*^(*x*_1_, *x*_2_) as a function of the species quantities *x*_1_ for the first cell and *x*_2_ for the second cell. In the figure, a vector field of white arrows depicts the net direction of probability flow due to the combined action of the internal auto-regulatory effects and the feedback control input *u*^*xAC*^(*x*_1_, *x*_2_) on the system. For practical implementation, the control input *u*′(*ϕ*) = *u*^*xAC*^(*x*_1_, *x*_2_) would be converted to light intensity through inversion of the calibration curve in Fig. 2B (blue line). The middle row of Fig. 4 shows the resulting steady state joint distribution of each condition, and in the bottom row of Fig. 4 shows the corresponding marginal distributions, with cell one in solid blue lines and cell two in dashed red lines. Fig. 4 shows that symmetry is broken only in the case where the cells’ genetic design includes auto-regulation *and* the feedback controller contains knowledge of the cells (i.e., the far right row of Fig. 4). In this best-case scenario, the two cells are very effectively driven each to their own unique and pre-chosen phenotype, irrespective of their initial conditions. If feedback is included without auto-regulation, the cells’ distributions are made tighter at some intermediate value between the target values for cell one and cell two; this results in a slightly better numerical cost value, but it becomes even less likely that both cells will reach their target phenotypes at the same time as compared to the uncontrolled situation (compare first and second row in Fig. 4). Conversely, if auto-regulation is included without feedback (i.e., the light level is fixed at some optimal value), the cells exhibit a bimodal distribution with some cells near each target value, but there is no means to control which cell expresses which phenotype, and the cost function is again worse than the case with no auto-regulation (compare first and third rows of Fig. 4). These data taken together suggest that auto-regulation and feedback control, in addition to intrinsic single-cell noise, are all critical to the break symmetry and enable differential control of multiples cells using a single input. Specifically, noise breaks the symmetry of cell behavior and allows cells to switch independently between phenotypes, feedback helps to reinforce this noise and steer cells toward desired phenotypes, and auto-regulation helps stabilize cell behaviors once they have attained their desired phenotypes.

**Figure 4:**
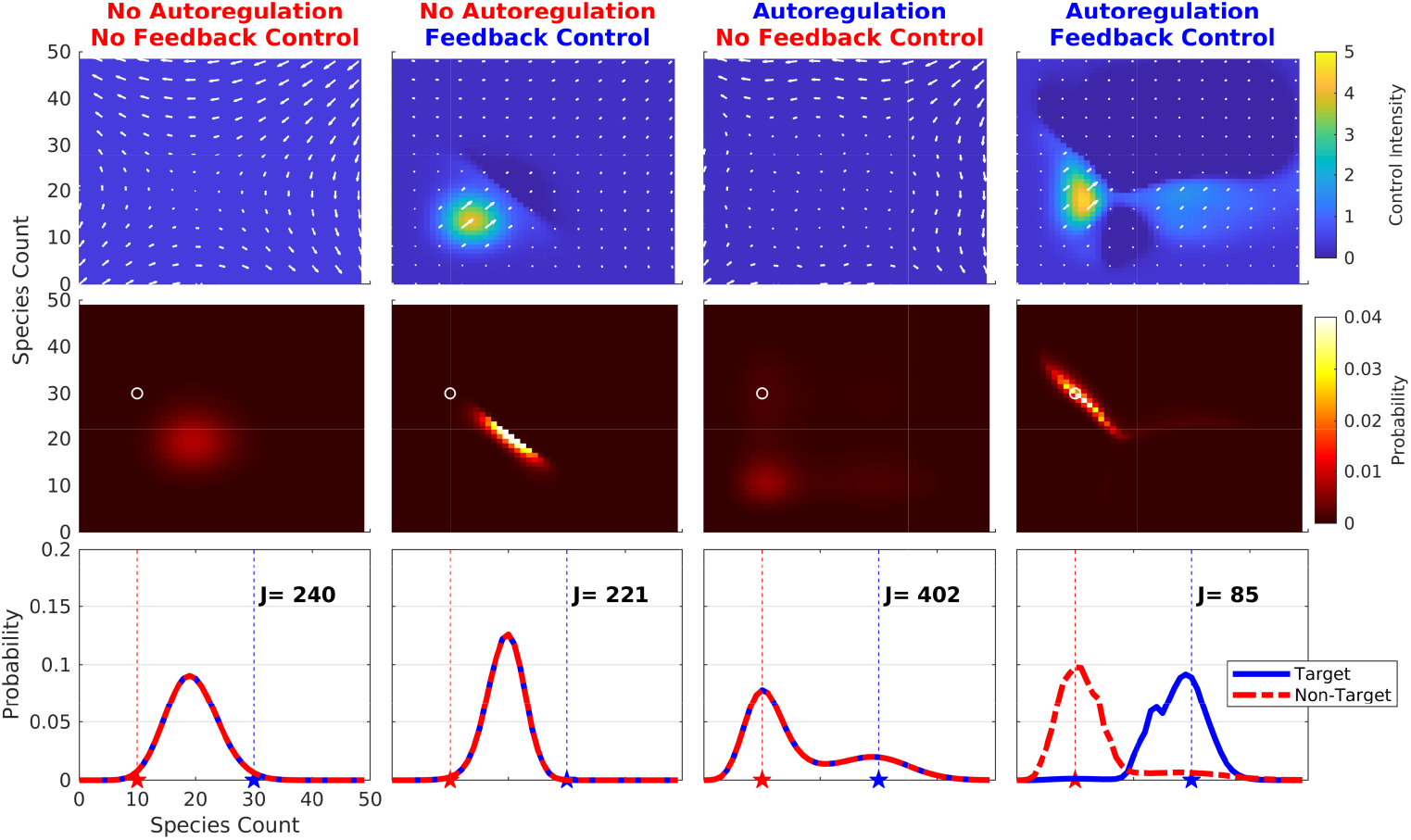
Optimized control laws and performance results for cells and different combinations of regulatory structure and control strategy. (Top row) Optimized control input versus cells’ states, **u**^*xAC*^(*x*_1_, *x*_2_) (colormap at far right). White vectors plotted over the control inputs represent the net flow of probability at that point in state space. (Middle row) Resulting joint probability distribution (**P**(*x*_1_, *x*_2_), colormap at far right). The target point *T* = [30, 10]^*T*^ is denoted by a white circle. (Bottom row) Corresponding marginal distributions for the cells **P**(*x*_1_) (blue) and **P**(*x*_2_) (red) and time-averaged cost function, *J*. Leftmost column shows results with no auto-regulation and a constant control signal. Second column shows the result with no auto-regulation and feedback control strategy. Third column shows the result with auto-regulation and constant control signal (UAC in text). Rightmost column shows the results for auto-regulation and fully aware feedback control (FAC in text).

### Effective differential control of many cells using a single input is possible, even when observations are limited to a single cell of interest

We next examined a more general problem to control an arbitrary number of cells simultaneously. In this case, we consider a situation where the controller acts on many cells simultaneously with the goal of steering a single observed cell (*x*_1_) to one state and all remaining unobserved cells to another state. To handle this problem, we first utilize a partially aware controller (PAC) that observes only 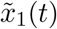 and ignores the states of all other cells. Fig. 5 compares the results of this simplified controller (middle column) to those of the two-cell controller from the previous section (left column). The resulting control law, *u*^*PAC*^(*x*_1_), is optimized to find the best control signal input for each possible value *x*_1_ of the observed cell, and the top row of Fig. 5 shows that this optimal PAC controller depends only on the observed cell (*x*_1_ axis) but is constant with respect to all unobserved cells (*x*_2_ axis). Despite the simplicity of this control strategy and the fact that it requires only knowledge of the instantaneous expression of the single observed cell, the second and third rows of Fig. 5 show that the PAC controller effectively breaks symmetry to force the observed cell to a high expression phenotype while most unobserved cells are correctly directed to the low expression phenotypes.

**Figure 5:**
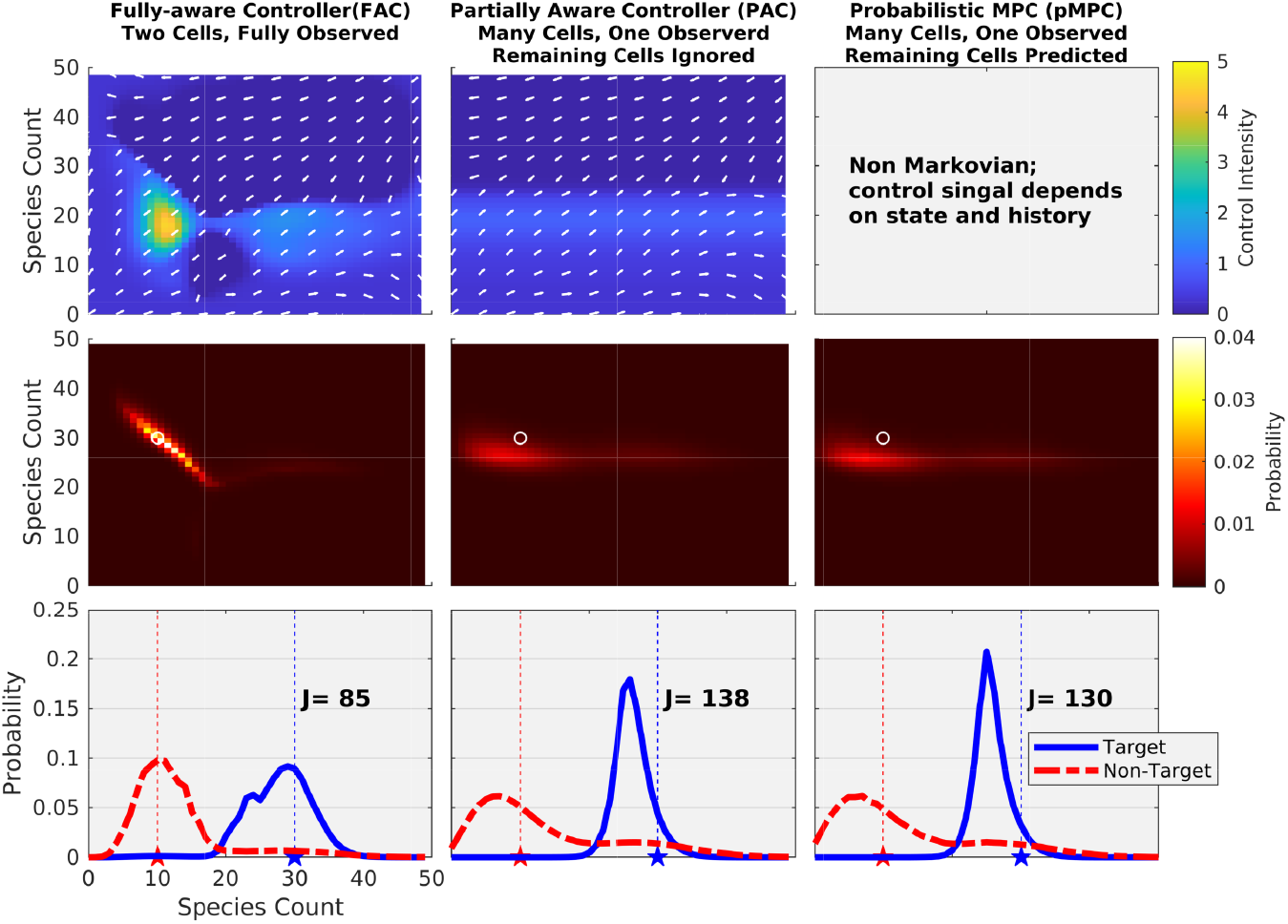
Control laws and performance for strategies to control many cells at once using partial knowledge only of a single observed cell. Each row shows: (top) the control law 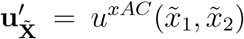, (middle) the resulting joint probability distribution (**P**(*x*_1_, *x*_2_)), and (bottom) the marginal distributions for the observed cell **P**(*x*_1_) (blue) and remaining cells **P**(*x*_2_) (red). The middle column shows results for the partially aware controller (PAC in text), where the control depends only on the single observed cell 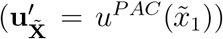. The rightmost row shows results for the probabilistic model predictive controller (pMPC), which also applies to an arbitrary number of cells. For the pMPC, the process is non-Markovian in that the controller 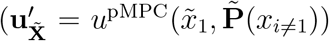 depends not only upon the state of observed 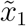, but also on the predicted probability vector for the unobserved cells, 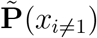. To enable comparison to previous two-cell cases, the leftmost column shows the results for the fully aware feedback control (FAC) with only two cells. Color bars are shown to the right of each row.

Although the partially aware controller under-performs compared to the fully aware controller for the case of exactly two cells (compare middle and left columns of Fig. 5), the advantage of the PAC is that it works equally well, and without any modification, for any arbitrarily large number of unobserved cells (Supplemental Fig. S1). In contrast, to use the FAC for more than two cells requires modification, such as training of a higher rank tensor representation of the control algorithm, or defining a control law based on the mean, median, or some other statistical quantity for the groups of cells to be assigned to each phenotype. Unfortunately, the former high-order tensor approach is computationally intractable using existing methods, and the latter approach rapidly loses performance as the number of cells is increased. For example, when the controller is based on the observation of 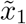 and the mean of the remaining cells 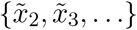 (FACM, see Supplemental Information), we observe that for any more than a single cell in the second group, the PAC outperforms the FACM (Supplemental Fig. S1, compare FACM and PAC controllers).

### A probabilistic Model Predictive Controller (pMPC) can improve the control of many cells using a single observer and a single input signal

In the previous section, the PAC control was based on only the observation of a single observed cell, and had no information about the other cells that it was also seeking to control. However, knowing the history of the input signal (i.e., the light intensity over time in the past), the FSP approach allows for the possibility that the controller can estimate the probability distribution of all non-observed cells. With this possibility in mind, we next explored a new class of controller that could use direct knowledge of the protein expression in the observed cell, the known control input signal at the current time, and a probabilistic model to predict the distribution of expression in the unmeasured cells. Specifically, we use the FSP approach to integrate our prediction of the probability distribution for the unobserved cells as:

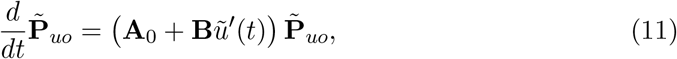

where 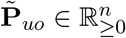 is the estimate of the probability mass vector for the protein expression in the unobserved cells, **A**_0_ ∈ ℝ^*n×n*^ is the infinitesimal generator for a single cell in the absence of any control input, and the scalar variable *ũ*′(*t*) > 0 is the instantaneous input signal that is produced by the controller. We note that the probability mass vector estimate 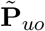 is the result of a stochastic process that depends upon the full history of the input signal *ũ*′(*t*).

Using this prediction for the unobserved cells, the pMPC controller law can now be defined as

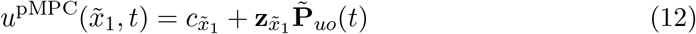

or written in vector form for all possible values of 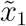 as:

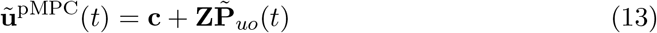

where **c** = [*c*_0_, *c*_1_, …, *c*_*n*−1_]^*T*^ is a constant vector in ℝ^*n*^, and **Z** ∈ ℝ^*n×n*^ is a matrix of linear weights which adjusts the input based off of the estimated unobserved probability distribution 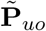. In our practical implementation, we assume that the controller in Eq. (13) is piecewise constant with respect to 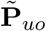 over a time step of 0.5 min, but it changes instantaneously with each even that affects the observed cell 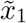. The weights of **c** and **Z** are then jointly optimized to minimize the cost function, *J*. The simple formulation of the control law in Eq. (13) admits the possibility for non-achievable negative values of light in order to construct a computationally tractable optimization procedure. However, in testing the controller, this non-physical situation is corrected by saturating negative control signals to zero in the true test of the system (see Supplemental Information). We note that this approximation to allow for negative control signals in the control law specification and subsequent correction to saturate these to zero in the control law test suggest that the pMPC controller identified here is sub-optimal. However, despite this non-optimal design, Fig. 5 shows that the resulting *non-optimized* pMPC (*J* =130) controller outperforms the *fully optimized* PAC (*J* =138), demonstrating that probabilistic model predictions can be used to improve control performance even in the absence of observations for many of the cells under its control. Having succeeded in our main goal to determine if probabilistic predictions could improve control results in principle, we leave the fine tuning of the specific pMPC control strategy to future investigations and more sophisticated control design strategies.

For a closer look at how the pMPC approach works to control observed and unobserved cells alike, Figs. 6A shows an example control input over time, and Fig. 6C shows the resulting trajectories over time for the observed cell (blue), the predicted probability distribution for unobserved cells (gray shading), and a representative unobserved cell (red). We reiterate that the controller has no direct knowledge of the red line. From the figure, it is clear that observed cell is well maintained near to its target value with low variability. Moreover, Fig. 6C shows that knowledge of the fluctuating input signal is sufficient to yield good predictions of the unobserved cell response (compare red line with dark gray shading), although as expected there are periods of poor predictions when the specific unobserved cell samples the higher or lower tail of the predicted distribution (e.g., at about 1900 minutes for the red curve in Fig. 6C). In addition to outperforming the PAC approach in terms of the overall cost function, the pMPC provides additional predictions for when the controller is effective, or when unobserved cells are more likely to escape from their intended phenotype. To illustrate this Figs. 6B shows the probability of observing fifteen or less protein molecules in the unobserved cell. When this probability exceeds a 90 percent threshold the control is expected to be effective, and the region is labeled as orange in Figs. 6B and Figs. 6C. Figs. 6D shows the marginal distributions averaged over all times, and Fig. 6E shows the marginal distribution only for the periods of time when the probabilistic model predicts its own effective control of the unobserved cells (orange regions). By focusing only on these times identified as successful by the controller, the cost function of the controller substantially decreases from *J* = 130 to *J* = 58. These results suggest that predicted dynamic information about the unobserved cells can not only be used to improve the quality of the controller, but that the pMPC can also be used to self-assess when control is working well, and when it is not.

**Figure 6:**
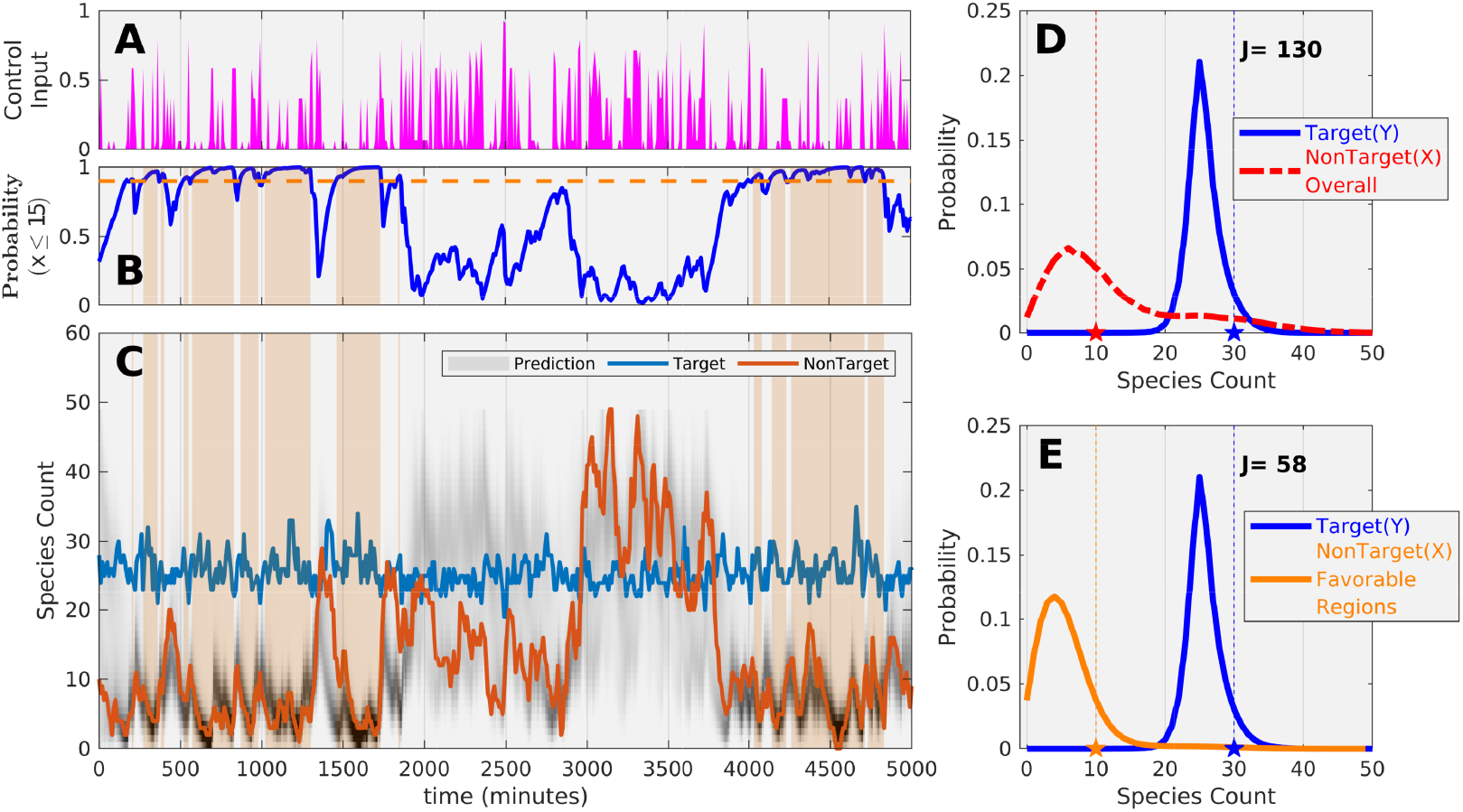
Control signal and response versus time for pMPC control. (A) Control signal generated by the pMPC controller. (B) Probability that unobserved cell is 15 or less versus time. When the probability is greater than 0.9 (orange line), then the system is considered to be in a “effective control” state. (C) Predicted transient distributions for unobserved cells (gray), observed cell (blue), and a single unobserved cell (red). Orange regions correspond to effective control times. (D) Time averaged performance of the control law in terms of marginal distributions for the observed call (blue) and unobserved cells (red). (E) Average of the pMPC performance, when considering only effective control periods.

### Controllers designed using simplified models can be effective to control more complicated processes with hidden mechanisms and dynamics

We next ask how well could controllers designed using simplified stochastic models work when they are applied to control more complex systems that contain additional hidden states and which have unknown dynamics or time delays. To perform this analysis, we first account for the difference in meaning and units for the input signal *u*′(*ϕ*) used in the reduced auto-regulation models (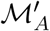 in Fig. 1C) and its analog *u*(*ϕ*) used in the full auto-regulation model ℳ_*A*_ in Fig. 1D). By using steady state ODE analyses of 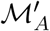 and ℳ_*A*_, we quantified the calibration curve to map inputs between the two models as shown in Fig. 7A. After calibration of the input signals, we verified that the full and reduced auto-regulation models result in similar bifurcation diagrams as shown in Fig. 7B. However, although calibration allows us to match both the quasisteady (i.e., very slow) and fast fluctuating input responses of the two models (Figs. 7C,E, respectively), the temporal responses to slow input frequencies are qualitative and quantitatively different, as can be observed by the different input-to output time lags and amplitudes in Fig. 7D.

**Figure 7:**
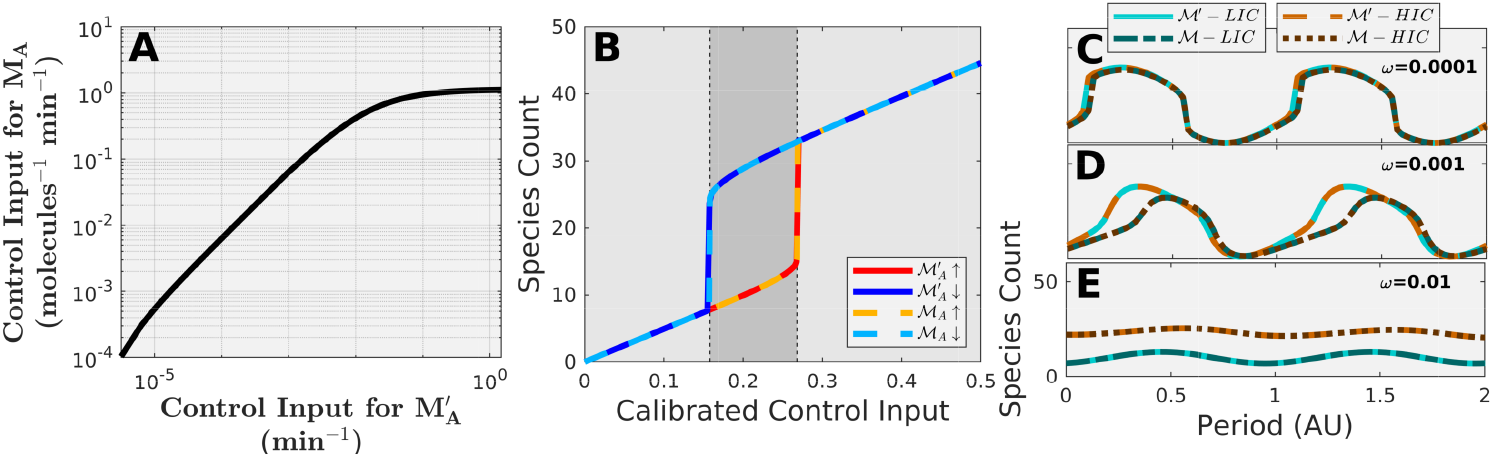
Calibration and use of controllers for use with a new, more complex model. (A) Calibration curve identified to match steady state ODE of simple and complex auto-regulation model (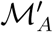 and ℳ_*A*_ from Fig. 1C,D). (B) Steady state analyses show that the calibrated inputs result in similar hysteresis behavior for both 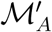 and ℳ_*A*_. (C) Input-output response analysis shows that ℳ_*A*_ with calibrated inputs closely matches behavior of 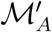 at ultra slow frequencies (0.0001 RPM), but (D) the more complex model begins to lag at slow frequencies (0.001 RPM). (E) At fast frequencies of 0.01 RPM, the complex auto-regulation model ℳ_*A*_ is able to retain memory of its initial conditions and again exhibits similar phenomena compared to the simplified model.

Having calibrated the controller for the full model to match the response of the simpler model, we then take the UAC, FAC, and PAC controllers from above and apply them directly (i.e., without any further tuning or optimization) to the full mechanistic model with auto-regulation. Despite the differences in temporal behaviors between the two models, the previously identified UAC and FAC controllers still work to break symmetry and drive both cells toward the correct differentiated phenotypes as shown in Fig. 8. We note that with further modifications, the control laws derived using the simplified model could certainly be improved for use in the more complex system. However, our primary goal was to explore how well designs made in one context should perform when used in another different context, and subsequent fine tuning for the complex model is left for future investigations.

**Figure 8:**
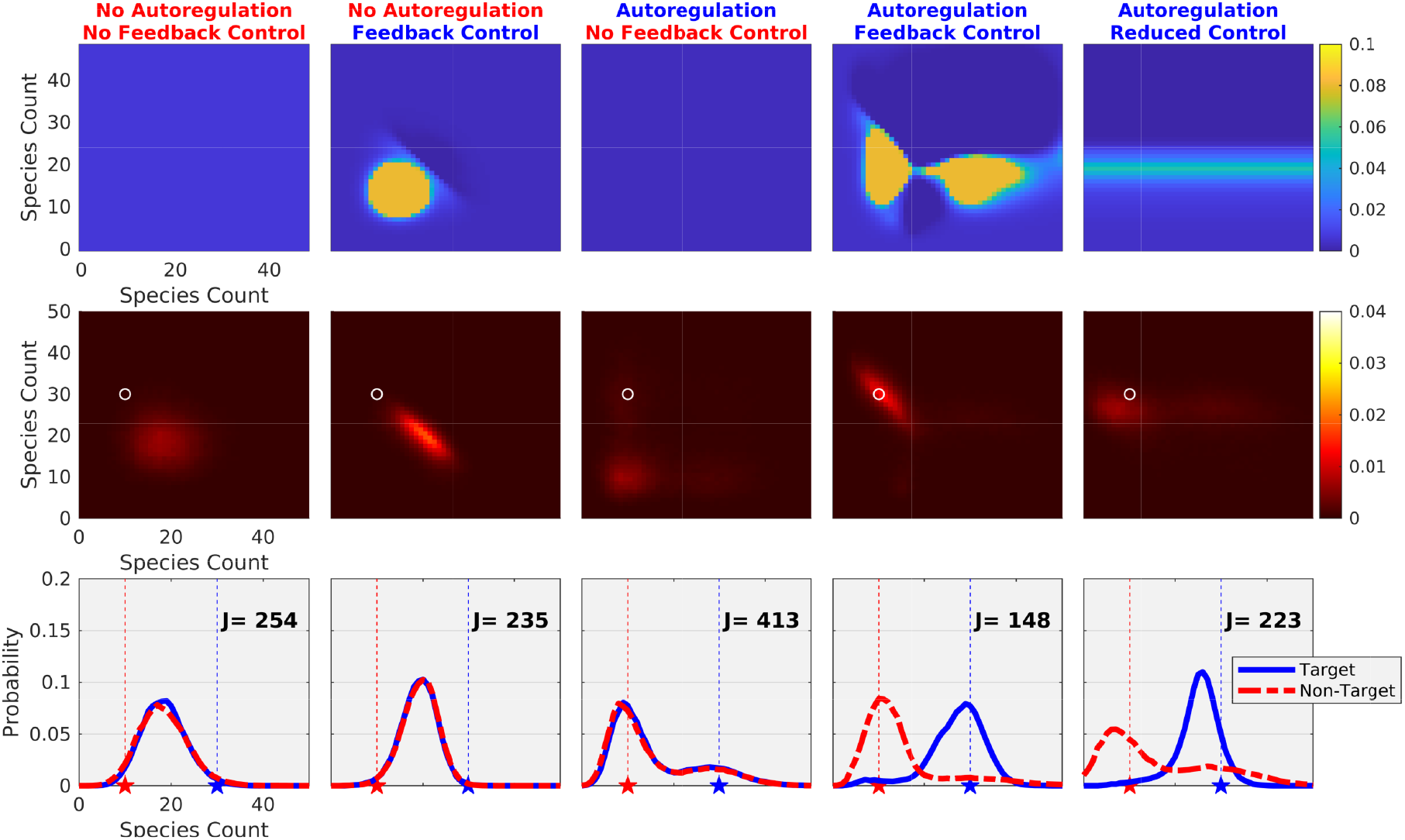
Full Models are paired with the calibrated controllers (top row) to solve for the joint probability distributions (middle row) and marginal probability distributions (bottom row). As before both auto-regulation and feedback are needed to break symmetry and control fails if either of these are missing (rightmost three columns). The fully aware controller (FAC, fourth column) successfully works to control two cells with the complex dynamics, and the partially aware controller (PAC, rightmost column) successfully can control a single observed cell to one phenotype and an arbitrary number of unobserved cells to another different phenotype.

## Conclusion

The treatment of noise in synthetic biology has largely been centered around the management of noise as a nuisance property that needs to be mitigated or eliminated. Despite improvements to minimizing noise in bio-circuits, noise largely remains a fundamental physical limit due to the combination of very small cell sizes, where single molecular events have increased importance, and increasing complexity of synthetic circuits, where most dynamical influences are unknown or unmeasured. The results here show how a few increasingly common synthetic biology motifs–such as optogenetic transcription factors, activatable polymerases and auto-regulation promoters–can in principle be combined to form regulatory modules and integrated with new external feedback controllers not only to mitigate intrinsic noise, but even to exploit that noise to achieve new multi-cellular behaviors.

Our work compliments that of previous efforts to develop single- and multiple-input-single-output (SISO and MISO) control for synthetic biology applications^40,41^ that have primarily sought to control one cell at a time or to control an entire population of cells to all reach the same phenotype. Specifically, we have demonstrated new types of optimizable single-input-multiple-output (SIMO) stochastic controllers that rely on the integration of noise, non-linear auto-regulation, and feedback to simultaneously control of multiple cells using a single chemical or optogenetic input. The first of these, the fully aware controller (FAC), assumes full knowledge of each individual cell’s behavior and achieves the best control performance. The disadvantage of the FAC is that it requires knowledge of each individual cell (i.e., optical tracking and image processing analyses) and computationally intensive operations both to solve multi-dimensional chemical master equations and to search very high dimensional spaces for optimal controllers. However, a second partially aware controller (PAC) requires only the knowledge of a single cell of interest, yet the PAC can control that cell to one phenotype and drive all others to an alternate phenotype with an accuracy almost equal to the FAC. The advantage of the PAC is that it is very easy to implement and optimize as the dimension of the control law must only consider the single observed cell. The third controller introduces a probabilistic model predictive control (pMPC) strategy that computationally predicts the probability distribution of all non-observed cells based on integration of the chemical master equation under the known history of the applied input signal. Although we envision that similar control strategies may have applications in other fields, such as for autonomous vehicle or smart grid applications, to our knowledge, the proposed pMPC approach is the first example of a hybrid control strategy that predicts and exploits noise and feedback to simultaneously and differentially control multiple identical agents using a single control signal.

In addition to noise, one of the most important challenges in model-driven synthetic biology, is that many important regulatory mechanisms are currently unknown, even for simple biological systems. Similarly, very few parameters are known at any level of certainty, and most of these parameters vary from cell to cell or situation to situation. Moreover, it is already extremely computationally expensive to combine all known mechanisms into a single computational model, and although such models can be useful to reproduce a variety of biological behaviors, ^51,52^ such whole cell models are far too inefficient to enable the vast numbers of different simulations needed to optimize a design or control strategy. To circumvent these concerns, we demonstrated how a highly simplified phenomenological model could be used to design a controller that could be easily recalibrated using steady state dose-response measurements and then applied directly to a control a more complex system with hidden dynamics and with qualitatively and quantitatively different dynamic response features. Although it is common practice to use simple deterministic models to guide engineering design of modern complex devices, this demonstration in the context of stochastic single-cell processes suggests that there is also hope for similar applications of simple models in synthetic biology.

Although the potential of our computational results remains to be verified through independent experimental investigation, we believe that this numerical demonstration of the potential for a new control paradigm not only opens new possibilities for integrated “cyber organic” approaches in synthetic biology, ^28–30,45^ but could also offer insight into natural cellular differentiation processes where cellular states are sensed, and control signals are transmitted, by neighboring cells. For example, it has been suggested that stochastic fluctuations in expression lead embryonic stem cells to achieve substantial, and functionally relevant heterogeneity in Nanog expression, where transiently low Nanog expression cells are prone toward differentiation, whereas high Nanog expression cells are less likely to differentiate. ^53^ As such, it might be interesting to explore the possibility that temporally controlled fluctuations in Nanog transcription factors ^54,55^ could selectively direct specific neighboring cells to differentiate while maintaining others in the stem state. Overall, we envision that advancing synthetic biology motifs, especially an increasing diversity of orthogonal transcription factors and promoters, ^12,56^ improved live cell reporters, ^13,57^ and faster and more specific optogenetically controlled transcription factors inputs,^30^ will integrate synergistically with new probabilistic model predictive control analyses to improve future efforts to understand how noise, non-linearity, and feedback combine to drive cell fate decisions in applications ranging from synthetic biofuel and biomaterial production to developmental dynamics or regenerative medicine.

## Methods

### Definition of Models in Terms of Stoichiometries and Propensities

To introduce our numerical approaches, consider a generic cell regulatory process that contains *N*′ distinct chemical species that interact with each other through *M*′ different reactions. At any point in time, the current state of the process in a *single* cell can be described by an *N*′-element vector **x** = [*x*_1_,…,*x*_*N*′_]^*T*^ ∈ 𝒳, where 𝒳 denotes the set of all possible states (e.g., the nonnegative spaces 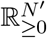 for a continuous process or 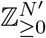 for a discrete process). The definition of the full state **X** for *multiple* cells is easily concatenated to consider a set of *N*_*c*_ individual cells

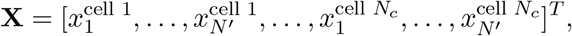

with an appropriate change to the total numbers of species (*N* ≡ *N*_*c*_*N*′) and reactions (*M* ≡ *N*_*c*_*M*′).

Under the assumption of a well-mixed spatial environment within each cell, one can define the dynamics of such a process by specifying the reaction stoichiometry vector and reaction rate for each *µ*^th^ reaction. ^58^ The *stoichiometry vector*, **s**_*µ*_ ∈ℤ^*N*^ is the net integer change in molecules after exactly one event of the *µ*^th^ chemical reaction (i.e., **s**_*µ*_ ≡ **X**(after *µ*^th^ reaction) − **X**(before *µ*^th^ reaction)). For continuous processes, the *reaction rate, f*_*µ*_(**X, Λ**, *u*) is a scalar that defines the speed at which the *µ*^th^ reaction would be expected to occur given the current state **X**(*t*), fixed physical parameters **Λ** and time- or state-varying control parameter *u*(**X**, *t*). For discrete stochastic chemical reactions, the reaction rate is replaced with a *propensity function w*_*µ*_(**X, Λ**, *u*)*dt*, which describes the probability that a single *µ*^th^ reaction would occur in the next infinitesimal time step of length *dt* given **X**(*t*), **Λ**, and *u*(**X**, *t*). For reduced order models 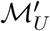 and 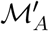, we replace *u* with *u*′ to denote the change in units needed for consistency with the model reduction.

### ODE Representation of Models

Using these simple definitions, one can easily write an ordinary differential equation (ODE) to define a deterministic description of the process dynamics as:

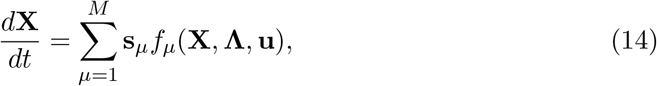

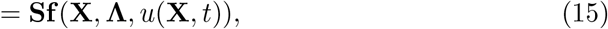

where **S** = [**s**_1_, …, **s**_*M*_] ∈ ℤ^*N ×M*^ is the stoichiometry matrix, and **f** (**X, Λ**, *u*(**X**, *t*)) = [*f*_1_, …, *f*_*M*_]^*T*^ ∈ ℝ^*M*^ *>* 0^*M*^ is the vector of reaction rates. For any given stoichiometry matrix **S**, and reaction rate function vector, **f** (**X**, Λ, *u*), the rate of change of **X** described by Eq. (14) can be integrated numerically to describe the system dynamics over time.

### Discrete Stochastic Representation of Models

For discrete stochastic systems, the specification of the reaction stoichiometry and propensity functions is sufficient to generate individual trajectories of the process using Gillespie’s Stochastic Simulation Algorithm (SSA, ^47^). Alternatively, one can also use these two properties to uniquely define the Chemical Master Equation (CME,^59^) as:

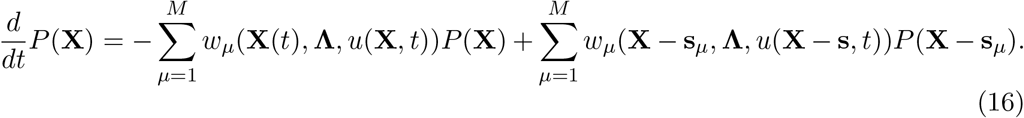

For this discrete state description, one can always enumerate all possible states as {**X**_1_, **X**_2_, …} ≡ 𝒳 and define a probability mass vector as the similarly ordered probabilities, **P** ≡ [*P* (**X**_1_), *P* (**X**_2_), …]^*T*^. Because the CME in Eq. (16) is linear in every term *P* (**X**_*i*_), it is often written in matrix format:

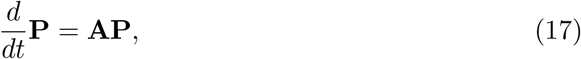

where the square matrix **A** is known as the *infinitesimal generator* and is defined directly from Eq. (16) as

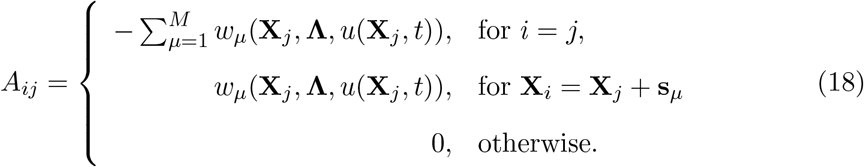

We note that the summation of *µ* = 1 to *M* = *N*_*c*_*M′* accounts for the increase in the number of possible reactions due to the existence of multiple cells. Because each term **X**_*i*_ refers to a specific enumerated state vector that is fixed in time, in the special case where *u* ≡ *u*(**X**_*i*_) (i.e., where *u* depends only on the current state and does not depend *explicitly* on time), the matrix **A** is constant with respect to time. For convenience, we can define the control parameter in vector form **u** = [*u*(**X**_1_), *u*(**X**_2_), …]. The final CME model with control can be written in simple form by separating the infinitesimal generator into it basal and control induced components as:

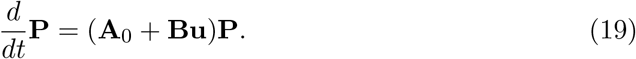

In this formulation, although the dynamics of each identical cell was independent and uncoupled in the basal infinitesimal generator **A**_0_, the added infinitesimal generator from the control input, **Bu**, can introduce coupling between cells. As an example, consider the fully aware controller (FAC) for two cells. The *i*^th^ state is written **X**_*i*_ = [*x*_*i*1_, *x*_*i*2_]^*T*^, and the control infinitesimal generator can be written as:

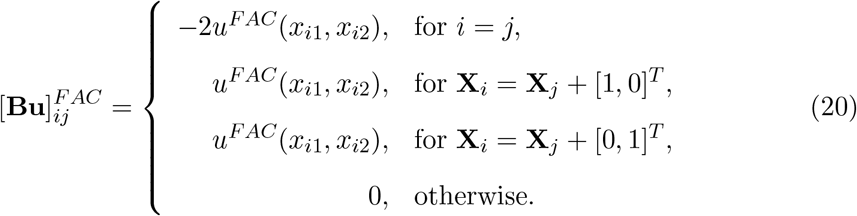

Similarly, for the partially aware controller (FAC) for two cells, the control infinitesimal generator can be written as:

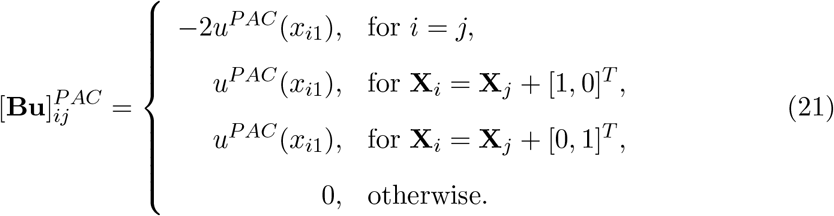

As discussed in the main text, in either case, the coupling introduced by the control infinitesimal generator [**Bu**]^*FAC*^ or [**Bu**]^*PAC*^ is sufficient to break symmetry and encourage cells toward desired differential expression phenotypes.

### Solution Scheme for Chemical Master Equation

To solve the CME in Eq. (19), we use the Finite State Projection (FSP^60^) approach, which truncates the allowable state space for every species and results in a finite dimensional ODE. However, it should be noted that the state space of a single arbitrary chemical species is given by the ordered set [0, 1, …, *n* − 1] up to some truncation limit *n*. The state space of multiple species, is enumerated by forming a tuple of all possible species available, each up to a similar maximum number. For a system of *N*_*c*_ cells with *N*′ chemically reacting species, where each species can range up to a maximum of *n* − 1 copies per cell, the number of distinct states is 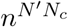, which quickly becomes intractable when *n, N*_*c*_, or *N*′ is large. For this reason, model reductions, simplifications, or approximations are essential, especially when these models are to be used with millions of different parameter sets when searching for optimal control strategies. Further, it is important to test and verify if control strategies designed and optimized using such simplified models will continue to be effective when applied to more general and more complex systems.

### Fitting of models to data

Fits of the reduced model to experimental data was performed numerically by optimizing both the set of model parameters and the calibration variables in unison. Since the parameter fits of 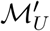 did not reveal a single set of unique parameters which fit data, the decay rate was calculated by hand by fitting the middle region of Fig. 2A and then fitting the parameters after fixing the protein decay rate. Fitting the ℳ_*U*_ to experimental data using calibration was performed by hand since mathematical tools to fit data often yielded poor results by becoming stuck in local minima. These hand fits were also constrained such that the decay rate of the protein is the same decay rate in 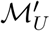.

## Supporting information

Supporting information is one file containing:

**Supporting Text** - supplemental details on feedback control designs.

**Supporting Figure S1** - Visualization of pMPC Control Law

**Supporting Figure S2** - Performance of stochastic controllers using varying numbers of non-target cells.

## Acknowledgments

Research reported in this publication was supported by the National Institute of General Medical Sciences of the National Institutes of Health under award number R35GM124747. The work reported here was partially supported by a National Science Foundation grant (DGE-1450032). Any opinions, findings, conclusions or recommendations expressed are those of the authors and do not necessarily reflect the views of the National Science Foundation. The funders had no role in study design, data collection and analysis, decision to publish, or preparation of the manuscript.

## Author Contribution

BM designed and acquired funding for the study. MPM and BM developed computational and theoretical tools. MPM performed all computational analyses. MPM wrote the manuscript. MPM and BM edited the manuscript.

## Data Availability

All data and computational codes needed to reproduce the figures in this manuscript are freely available through the GitHub page: https://github.com/MunskyGroup/Michael_May_et_al_2021

