## Supplemental Text and Figures for "Exploiting noise, non-linearity, and feedback for differential control of multiple synthetic cells with a single optogenetic input"

This supplemental information contains additional details about the specification and optimization of model-controller pairs as well as supplemental figures to support the minor results that are discussed in the main manuscript.

### Formulation and Optimization of Feedback Control Designs

The general tensor form of the FSP under state control can be written:

$$\frac{\partial P^i}{\partial t} = ([\mathbf{A}_0]^i_j + [\mathbf{Bu}]^i_j)P^j. \tag{22}$$

For the partial observation situation where not all states can be observed, we reformulate the FSP analysis as:

$$\frac{\partial P^i}{\partial t} = (A^i_j + B^i_{jm} M^m_\alpha v^\alpha)P^j, \tag{23}$$

where the indices $\{i, j, m\}$ refer to states in the Markov chain, and index $\alpha$ refers to the distinctly observable subset of those states that define the control inputs; $\mathbf{B}$ is the controller tensor as in the main text; and $\mathbf{v}$ the vector of control signals associated with each distinctly observable state. The control scheme used (i.e., FAC, PAC, or UAC) is defined $\mathbf{u} = \mathbf{Mv}$, where $\mathbf{M}$ is a lifting operator which takes the controller as defined on the observable state space $\mathbf{v}$ and lifts it into the proper dimensions to multiply with the control tensor. For the FAC control scheme, the number of elements in $\mathbf{v}$ matches the total number of states (i.e., $\mathbf{u} = \mathbf{v}$) meaning that $\mathbf{M}$ simply an identity matrix. For less than complete observation as in the PAC or UAC schemes, a single element of $\mathbf{v}$ can influence multiple parts of the state space, and $\mathbf{M}$ is a tall rectangular matrix that contains only ones or zeros, and $\mathbf{u} = \mathbf{Mv}$ describes the light activity in each state, while the lower dimensional $\mathbf{v}$ describes the smaller set of unique values of the control input for distinct observable states. With the UAC, the $\mathbf{M}$ matrix is represented by the $\mathbf{N} \times 1$ matrix filled with only ones and $\mathbf{v}$ is a scalar quantity.

Deriving how $\mathbf{P}$ changes with small changes in $\mathbf{v}$ at steady state gives

$$\frac{\partial \dot{P}^i}{\partial u^n} = 0^i_n = \partial_n((A^i_j + B^i_{jm} M^m_k v^k)P^j), \tag{24}$$

$$0^i_n = (B^i_{jm} M^m_n \delta^k_n)P^j + ((A^i_j + B^i_{jm} M^m_k v^k)P^j_n, \tag{25}$$

$$\frac{\partial P^j}{\partial u^n} = -[(A + BMv)^{-1}]^j_i B^i_{ko} M^o_n P^k. \tag{26}$$

Finally, plugging this into the definition for the objective score (Eq. 5 in main text), we have:

$$\frac{\partial J}{\partial v^n} = \frac{\partial (C_j P^j)}{\partial v^n} = C_j P^j_{,n} = -C_j [(A + BMv)^{-1}]^j_i B^i_{ko} M^o_n P^k. \tag{27}$$

With this expression in hand, $\mathbf{v}$ can be optimized by starting at an appropriate $\mathbf{P}$ and changing $\mathbf{v}$ in the direction of the negative gradient ($-J_{,n}$), and then updating the new $\mathbf{P}$. The process continues iteratively until convergence to a local minimum.

### Model Predictive Control

The probabilistic model predictive control (PMPC) uses a simple linear machine to generate light inputs, $\mathbf{u}(t)$, according to

$$\mathbf{u}(t) = \max\{\mathbf{0}, \mathbf{c} + \mathbf{Z}\mathbf{P}^{i}_{\text{nt}}(t)\}, \tag{28}$$

where $\tilde{\mathbf{P}}_{\text{nt}}$ denotes the random probability distribution of the unobserved cells given the history of $\mathbf{u}(\tau)$ for $\tau \in (0, t)$; the vector $\mathbf{c}$ provides the bias of the control signal with one entry for every possible value of the observed cell's value; and $\mathbf{Z}$ is a matrix which takes $\mathbf{P}_{\text{nt}}$ and outputs adjustments to the controller based on the probabilistic predictions for the unobserved cells. In this formulation, each row of $\mathbf{c}$ and $\mathbf{Z}$ represents the deterministic control bias and probabilistic correction for each state of the observed cell, given the history of the control signal. A heuristic optimization was performed on the system to tune the entries of $\mathbf{c}$ and $\mathbf{Z}$ and then simulating the process for long time periods. In circumstances where the proposed controller produces a non-physical negative control signals, the control signal is set to zero. Figure S1(A and B) shows the weights of $\mathbf{c}$ and $\mathbf{Z}$ after joint optimization was performed on the system.

### Distribution of Objective Scores

Analyses presented in the main text present the objective score, $J$ in terms of the expected Euclidean distance from the target state (Eq. 5 in main text). Certain controllers yield high variability in this distance over time in a single trajectory or if sampled for many different cells at a specific time point. To analyze this variability, SSA simulations were performed using each set of controllers and the score at each time point was measured. Histograms of score for time-series trajectories using the FAC, PAC, and pMPC show a long tail distribution of the score (Fig. S1C). These data taken together suggest that the score is often much lower than the expected value, but the average performance, $J$, is dominated by rare moments in time where the performance is poor, causing a large temporary increase in score and raising the average despite overall good performance. Small changes to the highly weighted tails can yield better average scores while returning remarkably similar distributions.

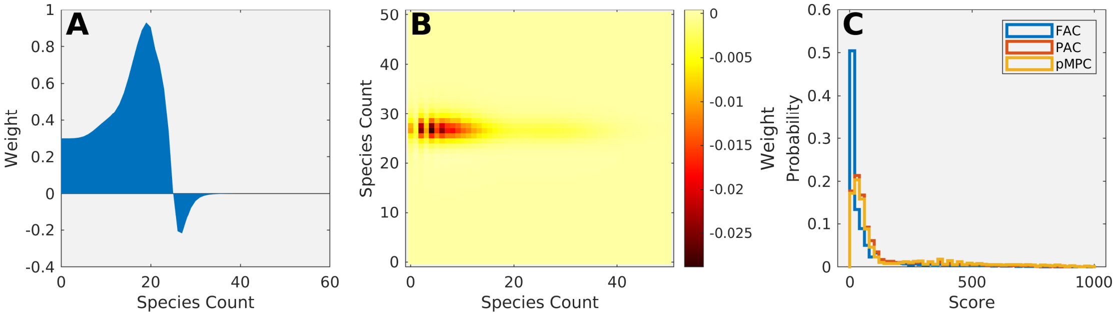


Figure S1: **Visualization of pMPC Control Law.** (A) Weights of $\mathbf{c}$ show that the pMPC tends to increase the controller when the observed cell is below 20, but tends to decrease the control signal when $\mathbf{P}_n t$ is weighted above 20 and turns off when the observed cell is above 30. (Z) Weights of Z show that the pMPC optimization preferred to weight the control input down when both the observed cell was above twenty five and the unobserved cell was near ten. (C) Distribution of scores obtained during time trajectory show that score over time is a heavy tailed distribution. Although the probability of a high score is low, the score value itself tends to be vary large and can increase variability in simulations as well as attributing large differences in score for similar looking distributions.

### Quantification of Controller Performance for Multiple Cells

We considered a set of five controllers, including the FAC, PAC, and pMPC controllers already described in the main text in addition to two adhoc controller that use (i) the mean of the FAC controller over all cells to be driven to the second target state (MFAC):

$$u_{\mathrm{MFAC}}(\{x_1, x_2, \ldots\}, x_N) =< u_{\mathrm{FAC}}(x_1, x_{i>1}) >, \tag{29}$$

and (2) the FAC controller evaluated at the mean of all unobserved cells (FACM):

$$u_{\mathrm{FACM}}(\{x_1, x_2, \ldots\}, x_N) = u_{\mathrm{FAC}}(x_1, < x_{i>1} >). \tag{30}$$

To quantify the performance of each controller under varying numbers of cells, we generated 32 independent simulations over 99,000 minutes (following a 1,000 minute burn in period) while sampling every 100 minutes using two, four, eight, and sixteen cells in the second group. The objective score for each simulation was computed by averaging $J$ over its corresponding trajectory. The lines depicted shown in Fig. S2A show the median of the 32 independent objective scores (colored line) as well as the 25th and 75th percentile (shaded regions) versus the number of cells. These data in Fig. S2 show that the performance of the FAC, MFAC, and FACM outperform the UAC, but this performance decreases as the number of cells increases. However, the PAC and pMPC performance is independent from the number of cells. These data taken together suggest that the PAC and pMPC are both better controllers once the number of cells is greater than or equal to two, and attempts to extend the FAC to work on summary statistics from many cells does not appear to be effective. Improved FAC controllers would be possible using higher order tensor control formulations (as opposed to summary statistics described here), but the computational complexity for the design of such control algorithms is currently prohibitive and left to future work.

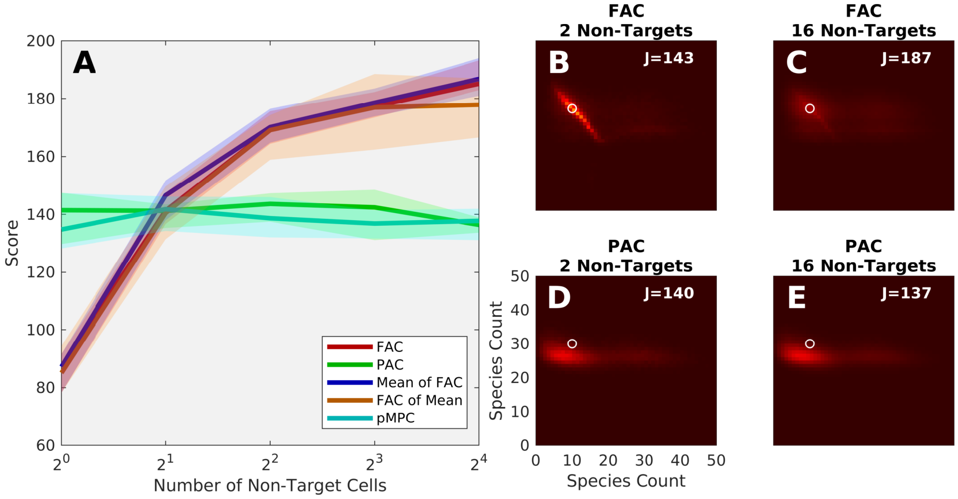


Figure S2: **Performance of stochastic controllers using varying numbers of cells.** (A) median score of 32 simulations using a set of five controllers in colored lines, with 25% and 75% quartiles shown in the color-shaded region. (B and C) FAC controller joint distribution of two cells chosen from a set of two (B) or sixteen (C) cells total, shows rapid degradation of performance when more cells are considered. (D and E) PAC controller joint distribution of two cells chosen from a set of two (D) or sixteen (E) cells total shows no change in the joint distribution.
